# Fungal Feature Tracker (FFT): A tool for quantitatively characterizing the morphology and growth of filamentous fungi

**DOI:** 10.1101/659672

**Authors:** Guillermo Vidal-Diez de Ulzurrun, Tsung-Yu Huang, Ching-Wen Chang, Hung-Che Lin, Yen-Ping Hsueh

## Abstract

Filamentous fungi are ubiquitous in nature and serve as important biological models in various scientific fields including genetics, cell biology, ecology, evolution, and chemistry. A significant obstacle in studying filamentous fungi is the lack of tools for characterizing their growth and morphology in an efficient and quantitative manner. Consequently, assessments of the growth of filamentous fungi are often subjective and imprecise. In order to remedy this problem, we developed Fungal Feature Tracker (FFT), a user-friendly software comprised of different image analysis tools to automatically quantify different fungal characteristics, such as spore number, spore morphology, and measurements of total length, number of hyphal tips and the area covered by the mycelium. In addition, FFT can recognize and quantify specialized structures such as the traps generated by nematode-trapping fungi, which could be tuned to quantify other distinctive fungal structures in different fungi. We present a detailed characterization and comparison of a few fungal species as a case study to demonstrate the capabilities and potential of our software. Using FFT, we were able to quantify various features at strain and species level, such as mycelial growth over time and the length and width of spores, which would be difficult to track using classical approaches. In summary, FFT is a powerful tool that enables quantitative measurements of fungal features and growth, allowing objective and precise characterization of fungal phenotypes.

**Author Summary:** One of the main obstacles to study filamentous fungi is the lack of tools for characterizing fungal phenotypes in an efficient and quantitative manner. Assessment of cell growth and numbers rely on tedious manual techniques that often result in subjective and imprecise measurements. In response to those limitations, we developed Fungal Feature Tracker (FFT), a user-friendly software that allows researchers to characterize different phenotypic features of filamentous fungi such as sporulation, spore morphology and mycelial growth. In addition, FFT can recognize and quantify other fungal structures including the fungal traps developed by nematode-trapping fungi. In order to show the capabilities and potential of our software, we conducted a detailed characterization and comparison of different fungal species. Our comparison relies on a series of experimental set-ups using standard and easily accessible equipment to ensure reproducibility in other laboratories. In summary, FFT is an easy to use and powerful tool that can quantitatively characterize fungal morphology, cell number and quantitatively measures the filamentous growth, which will allow advance our understanding of the growth and biology of filamentous fungi.

## Introduction

Filamentous fungi are among the most ubiquitous organisms on Earth. Their unique way of growth, i.e., by developing complex networks, allows them to survive in and colonize even the most inhospitable environments (1). Filamentous fungi can easily adapt to different environmental conditions (2) and change their nutritional requirements according to resource availability (3). One of the best examples of such adaptation is nematode-trapping fungi (NTF), which develop complex trapping devices under low nutrient conditions to capture and consume nematodes (4). Consequently, these fungi have great potential as biocontrol agents against parasitic nematodes (5). However, they often fail to establish in agricultural soils and their trapping behavior is rarely observed under natural conditions (6). Therefore, in order to employ NTF to control nematode populations, more detailed characterization of their growth and trapping behavior and a deeper understanding of their biology is urgently needed (3). One of the main obstacles to studying these fungi, and filamentous fungi in general, is the lack of tools for efficiently characterizing their growth and morphology (7,8).

The growth of filamentous fungi is often characterized using measures such as colony radius or fresh/dry weight (9–14), which do not capture the complexity of the mycelium. Other features such as conidiation or spore formation are frequently measured by means of time-consuming manual techniques (11,13,15). Moreover, other fungal structures such as the traps developed by NTF are measured manually (10,12,16). Thus, it is challenging to produce objective and quantitative results, and to capture subtle phenotypic differences using these approaches. In addition, vague and subjective terms such as “irregular growth”, “bulbous” or “fluffy” are commonly used in the literature to describe the morphology of fungal colonies (17,18). These imprecise descriptions make comparison of growth phenotypes between different mutants difficult and represent a limitation for the study of fungal biology.

Fortunately, fungi grow at a spatial and time scale that can be easily captured using basic imaging devices. Thus, together with the increasing availability of new technologies, this feature has resulted in the development of image analysis techniques to characterize different aspects of fungal biology (19), including programs to characterize germination of fungal spores (20), differentiation of aerial spore types (21), and measurement of the area of fungal cultures (22). However, these tools are often developed for very specific experimental setups and require prior image analysis or programming experience, limiting their accessibility to fungal biologists.

In response to these challenges, we have developed Fungal Feature Tracker (FFT). This user-friendly tool automatically characterizes several fungal phenotypes using images that can be obtained by means of basic imaging devices. FFT can be used to study different fungal species and conditions at various image scales and resolutions. Relying on simple built-in image analysis functions, FFT is able to quantitatively characterize conidiation, conidia morphology and different aspects of the mycelium, including the growth area, number of hyphal tips and total length of the hyphae. In addition, we have developed a function within FFT for detecting and counting the traps developed by NTF.

Here, we introduce FFT to the fungal research community. To illustrate the potential of this tool, we conducted a detailed phenotypic comparison of different fungal species and strains. We present a set of simple experimental set-ups that rely on basic methods and devices that can be used in most laboratories to characterize fungal phenotypes. The results of our phenotypic analyses demonstrate that FFT detects subtle differences in morphology and growth at both strain and species level. In summary, we believe that FFT will prove to be a valuable tool for mycology labs to automatize daily tasks and produce quantitative data, likely generating novel findings and providing additional insights into fungal biology.

## Results

### Quantitative measurements of conidia morphology

The spore morphology function of FFT detects different sizes and shapes of spores and conidia, and measures their different morphological features. In order to illustrate the power of FFT, we compared the conidial morphologies of five different filamentous fungal species (*Arthrobotrys oligospora* (*TWF154*), *A. musiformis, A. thaumasia, Trichoderma reesei*, and *Neurospora crassa*) in terms of their length, width, area and circularity. For each fungal species, we tested different parameter combinations in the calibration tab and determined the final parameter values by visual examination of the results. Subsequently, we used this parameter configuration to execute the spore morphology function of FFT on 10 images displaying between 1 and 3 conidia. FFT accurately detected the conidia of all studied fungal species despite their clear differences in shape and size (Fig. 1, top; note how FFT correctly delineated the conidia in blue). In addition, the program can be used to measure length and width (blue lines), given by the longest and shortest axes of the best-fit ellipse, respectively. These measurements obtained for each single conidium are shown in the bottom panels of Fig. 1.

**Figure 1.**
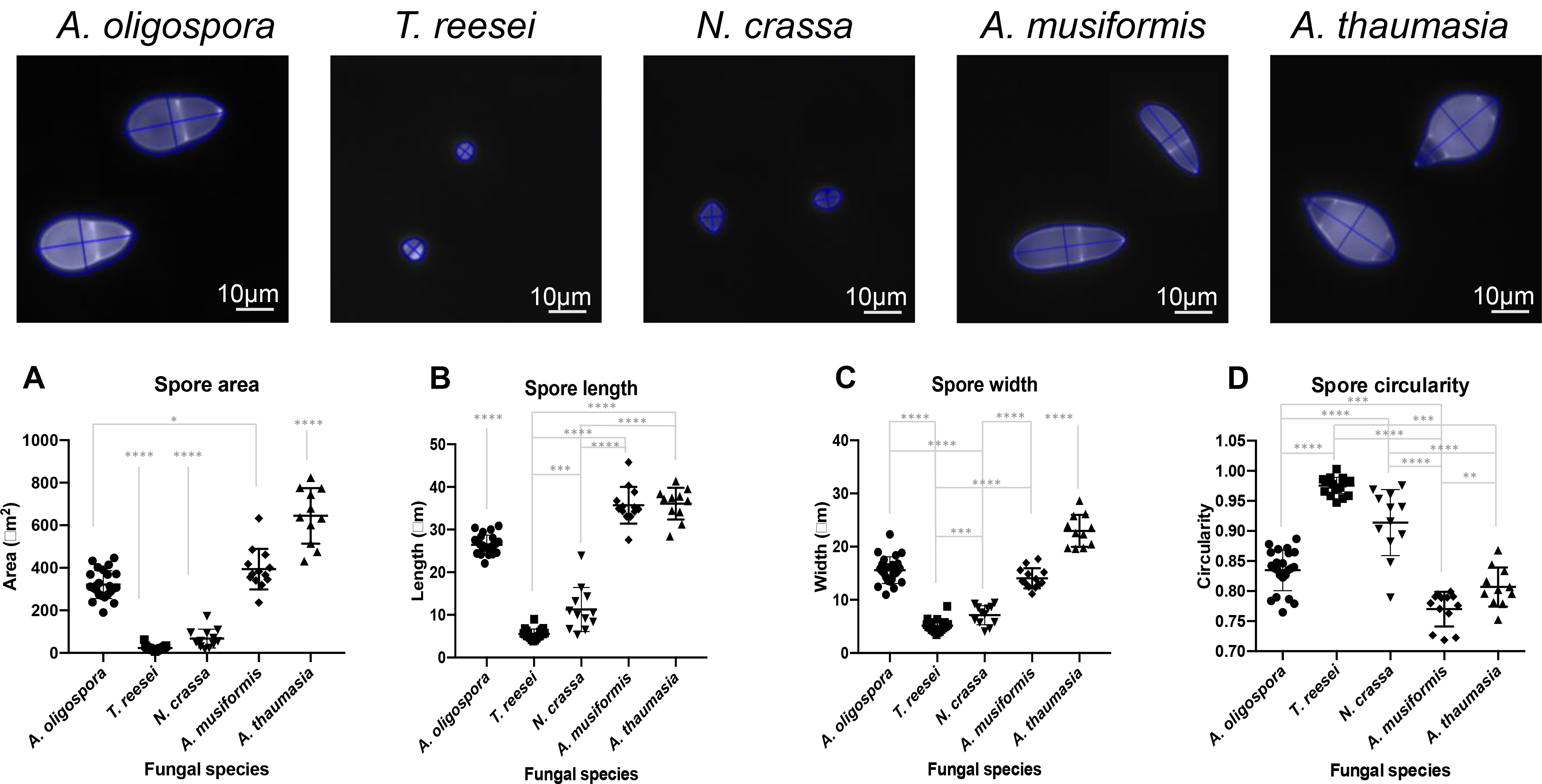
Characterization of the conidial morphology of five different fungal species using FFT. Results generated by the conidia/spore morphology function of FFT for images representing conidia of five different species (Top). Area (A), length (B), width (C) and circularity (D) of conidia for different fungal species, as assessed from 10 images per fungal species by FFT.

Conidial area varied significantly among the different fungal species (Fig. 1A). Moreover, FFT detected a substantial difference in the size of conidia from NTF, which were bigger than those of the other assessed fungi. For instance, *A. thaumasia* presented the largest conidia (average size ∼645 μm^2^), whereas the smallest conidia were those of *T. reesei* (average size ∼22 μm^2^). The results obtained for conidial length and width (Fig. 1B and 1C) are consistent with the data on conidial area. Within the NTF, *A. musiformis* conidia were of similar average length to those of *A. thaumasia*, whereas average conidial width in *A. musiformis* was closer to that of *A. oligospora*, resulting in values for *A. musiformis* conidial area lying between those of the other two species.

To measure conidial shape, FFT computes circularity (Fig. 1D). Circularity values close to one, such as those observed for *T. reesei* and *N. crassa*, correspond to almost perfect circles. This feature could be clearly observed for the detected conidia (Fig. 1, top), being very rounded for *T. reesei* and *N. crassa* and elongated in the nematode-trapping species.

In addition, we employed FFT to capture morphological differences among the conidia of three strains of *A. oligospora* (Supplementary Fig. S1). However, very few differences were observed for the morphological features we assessed. Only the length of the conidia produced by strain TWF132 was slightly longer than that of the other two strains.

### Automated spore counting by FFT

Conidia quantification is a daily task in many fungal laboratories. Consequently, we assessed the versatility of the spore/conidia counting function of FFT using a simple experimental set-up. For this purpose, we tested FFT using images captured at different resolutions of the fungal species and strains described in the previous section, thus representing a wide range of conidial sizes and shapes. When acquiring these images, it was important to take into account magnification and resolution since spores/conidia should be represented by more than one pixel in order to be efficiently detected. For example, images obtained with a basic digital camera and a microscope up to a magnification of 16X and average resolution will produce acceptable results when studying large spores such as those produced by NTF. In contrast, images obtained at the same magnification of *T. reesei* or *N. crassa* conidia may contain small dust particles or lighter areas that could give rise to false positive data (noise detected as conidia).

Figure 2 shows the conidia detected and computed by FFT from images obtained at 80X magnification (image size ∼ 1.3 mm x 1 mm). FFT excluded hyphae and recognized conidia of different size and shape, including the small conidia of *T. reesei* and *N. crassa*. The performance of FFT’s spore/conidia counting algorithm in terms of percent error is also shown in Fig. 2. To compute performance, we manually counted the number of conidia in 10 images per species and compared this number to that obtained using FFT on the same images using the following formula:

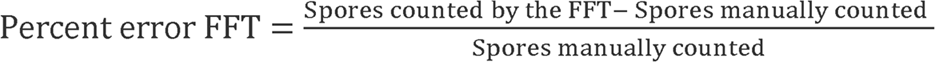

**Figure 2.**
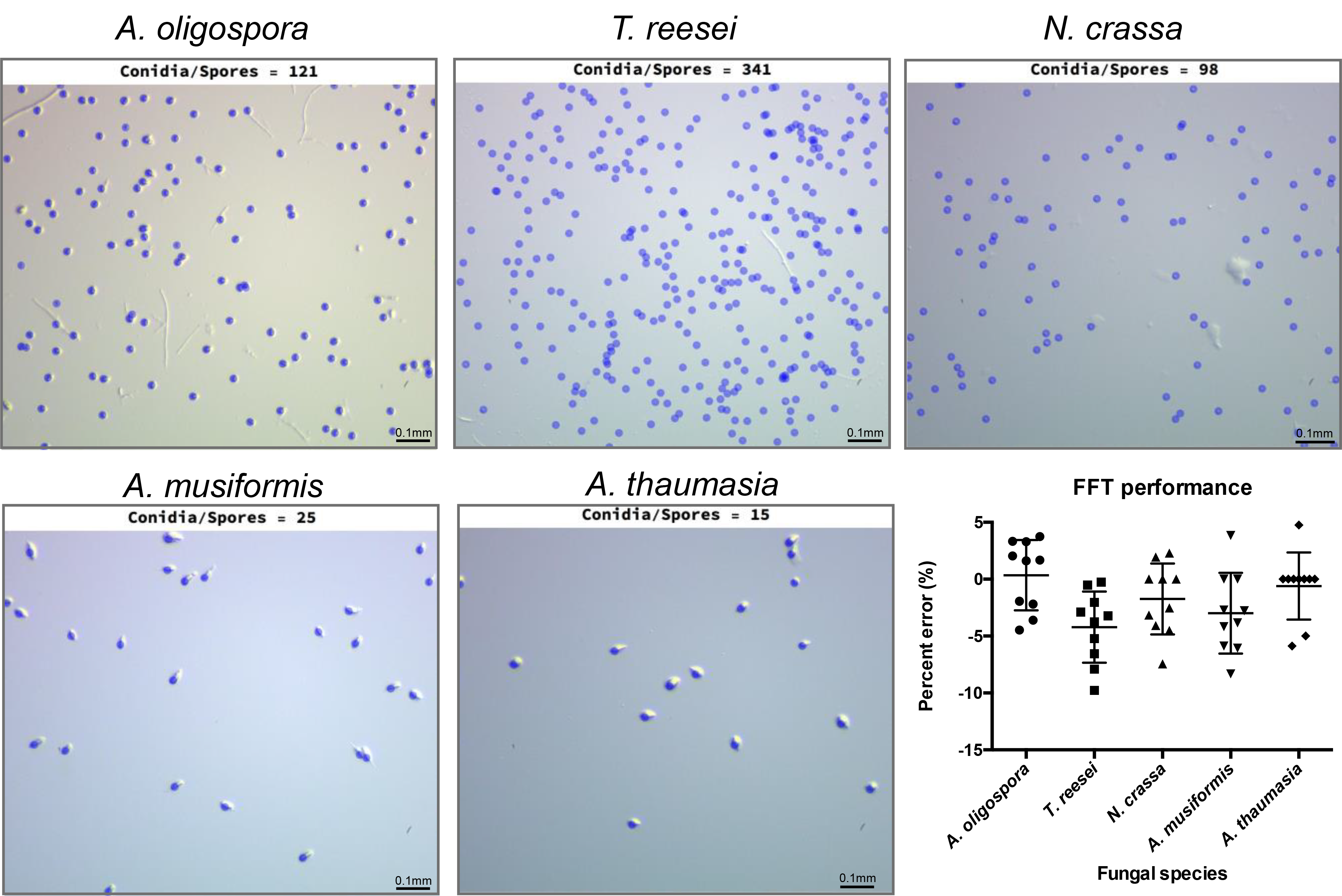
Results of the conidia/spore counting function of FFT for five different fungal species. Example of the conidia detected by FFT (blue) overlaid on the original images for each of the studied fungal species. Bar chart representing the performance of FFT as percent error between the conidia detected by FFT versus the number of manually counted conidia for 10-15 images per species.

Using this measure, percent error is positive when FFT detects more spores that when manually counted and is negative when FFT misses spores that where manually counted. We observed that average values of percent error are close to 0 for most species, which indicates a good overall performance of FFT in terms of conidial counts. Even though we used the same parameters (see the section “**Fungal Feature Tracker (FFT) interface and workflow**” below) to compute the number of conidia in all images of each of the fungal species we assessed, the percent error is below 5% for most of these images. The spores of *T. reesei* are small and numerous (average of 380 conidia per image), meaning that noise in the form of dust or irregularities in the media can be easily confused with small conidia. Therefore, we adopted a conservative approach and selected restrictive parameters to ensure that only conidia clearly distinguishable as such are detected by FFT. This approach results in a slight underestimation of the numbers of *T. reesei* conidia, as shown in Figure 2.

We also studied conidiation in different strains of *A. oligospora* after 7 days of growth. Since *A. oligospora* conidia are larger than those of other fungi (Fig. 1), we could detect them at a magnification of 16X. Images obtained at that magnification contained all the conidia in a 5 μl drop of spore solution. Therefore, we were able to compute the total number of conidia produced by each strain in an entire 5 cm Petri-dish after 7 days of cultivation. Figure 3 shows the results obtained by FFT for 10 images per strain. Strain TWF154 produced the most conidia, with an average of 848 conidia per 5ul of spore solution, equivalent to 169,600 conidia per Petri-dish, whereas strain TWF132 produced the least amount of conidia per Petri-dish (Fig. 3).

**Figure 3.**
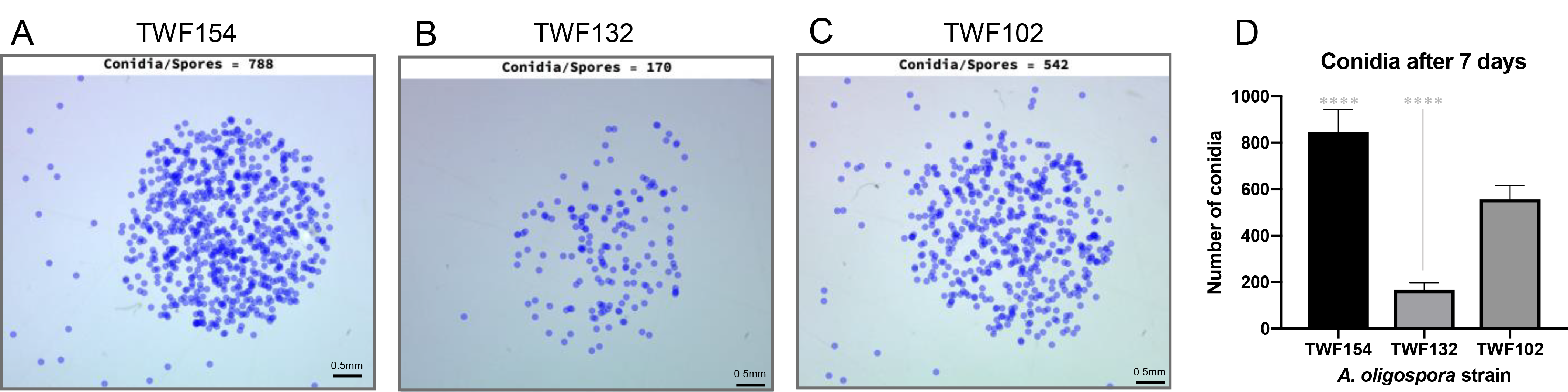
Differences in conidiation for three strains of *A. oligospora* as evaluated by FFT. Number of spores detected in a 5 μl droplet of spore solution extracted from cultures of strains TWF154, TWF132 and TWF102, respectively (A-C), together with a bar chart representing the results obtained from 10 droplets per strain (D).

### Comparison of mycelial development among different fungal strains, species and media conditions using FFT

The mycelium characterization function of FFT quantitatively computes several fungal measures from images of the mycelium such as total length, area covered by the mycelium, and the number of hyphal tips. This function works both on single images and on sets of images representing a temporal series, thus reflecting mycelial growth.

In order to achieve high quality results, the mycelium characterization function of FFT requires high contrast images with low noise. Such images can easily be obtained using fluorescent dyes to generate images showing a bright mycelium on a dark background so that noise (such as media irregularities) is not observed. Several fluorescence dyes require cell fixation or can stain mycelia for only short time periods (23–25). However, SCRI Renaissance 2200 (SR2200) can stain fungal cell walls without arresting growth (26), allowing mycelial development to be captured in high quality images at consecutive time-points.

We tested the accuracy of the mycelium characterization function of FFT using a set of 10 fluorescence images showing different stages of the mycelium developed by *A. oligospora* (strain TWF154). We constructed ground truths of each image and computed precision, recall, F-measure and Matthews Correlation Coefficient (MCC), all of which are measures widely used to assess the performance of image detection algorithms (27). In Table 1, we present the values obtained for each of these measures for the mycelium detected by FFT from each of the 10 images. For all these measures, a value of 1 represents a perfect detection, so values close to 1 represent accurate detections. The averages for all the studied measures are high, indicating that the images detected by FFT accurately represent the mycelia.

**Table 1.**
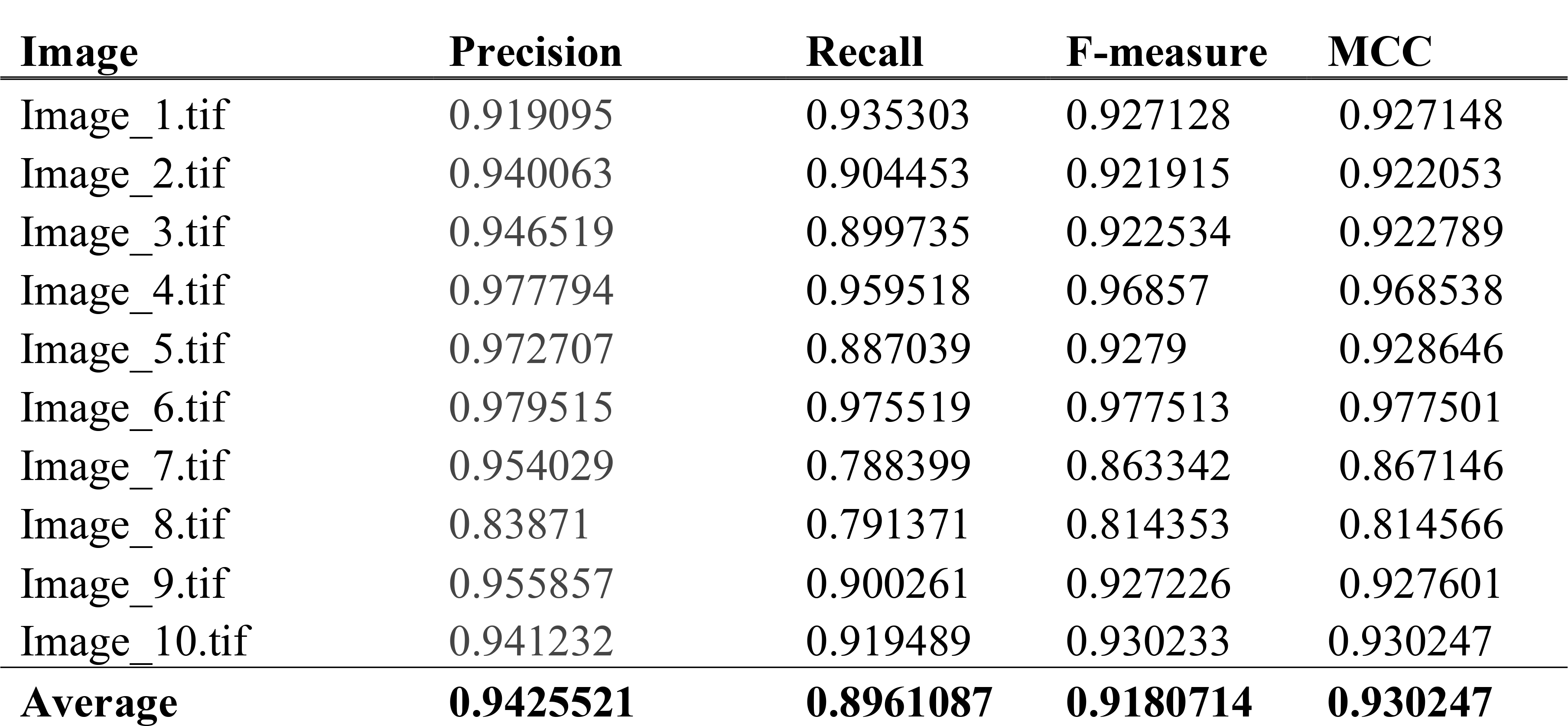
Accuracy measures of FFT analysis of 10 images of *A. oligospora.* Precision, Recall, F-measure and MCC computed for binary images assessed using FFT with respect to its ground truth counterpart for each image of our test set.

As an example, we show in Figures 4A-C the changes in mycelia of a stained *A. oligospora*, as tracked by FFT from a time-series of images. Figures 4A-C represent composites of data obtained at each time-point, including the number and positions of hyphal tips (A), the area covered by the mycelium over time (calculated as the convex hull of the mycelium) (B), and the extent of the mycelium at each time point (presented as a colored heatmap) (C), with this latter revealing how total mycelial length (computed as the length of all hyphal segments) changes over time (see the section “**Mycelial characterization**” under “**FFT Functions**” below for details of how these characteristics were defined).

**Figure 4.**
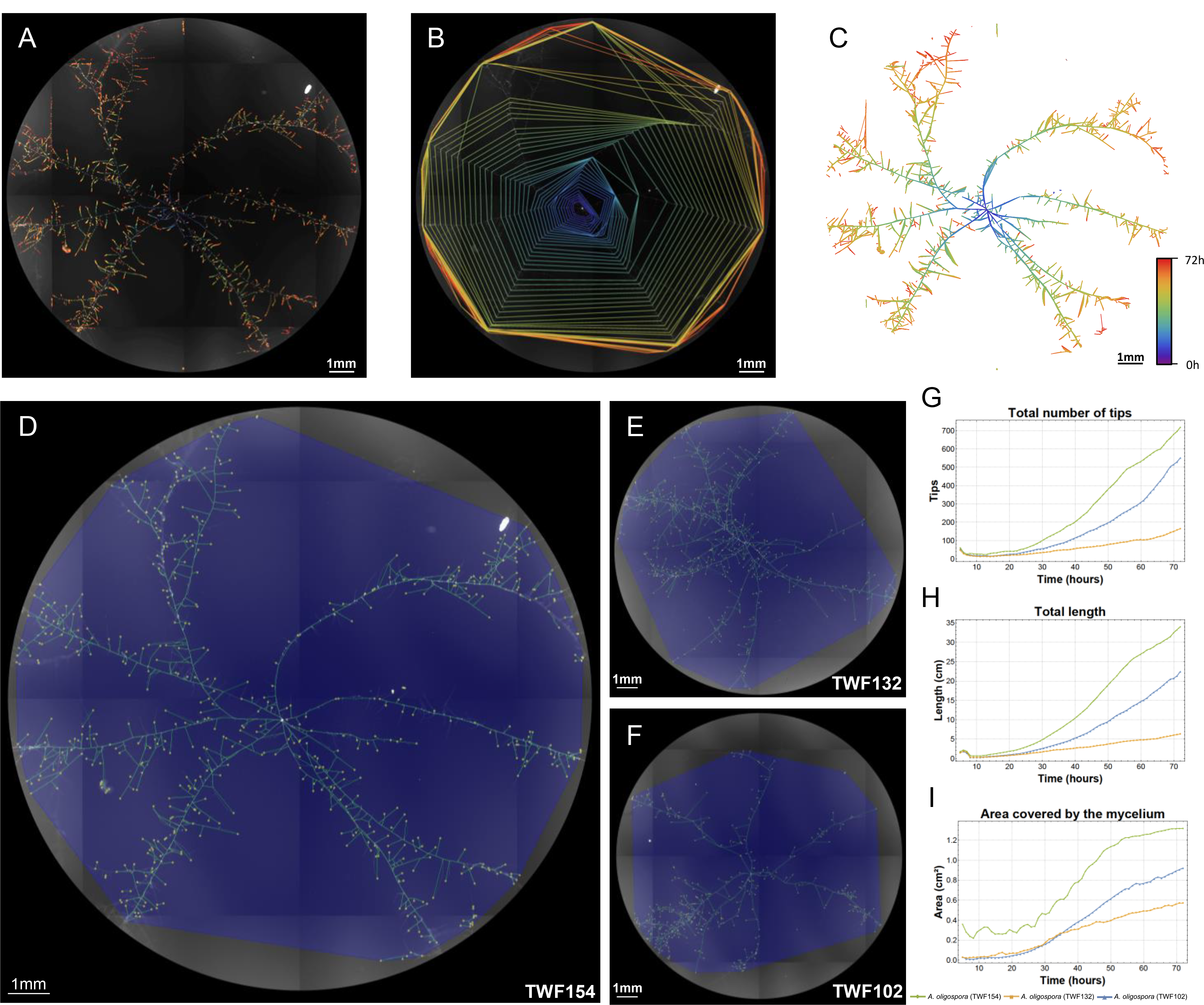
Output of the mycelium characterization function of FFT for different *A. oligospora* strains. Changes in the developing mycelium of *A. oligospora* (strain TWF154) over 72 hours, computed from the time-series of mathematical graphs generated by FFT (Top). A. Change over time of the area covered by mycelium, computed as the convex hull of all nodes in the mathematical graph for each time-point. B. Positions of the hyphal tips at each time-point. C. Overlay of the hyphal segments (edges) at different time-points showing development of the mathematical graph over time. Fungal features detected in images of the TWF154, TWF132 and TWF102 strains after 72 hours of growth (D, E, F, respectively). Area covered by the mycelium (blue polygon), mathematical graph representing the hyphal segments (green network), and hyphal tips (yellow dots) detected by FFT. The graphical representations show changes over time for the number of hyphal tips (G), total mycelial length (H), and the area covered by the mycelium (I), computed as the mean of six replicates per strain and time-point.

We studied growth from single spores on LNM over 72 hours for three different strains of *A. oligospora* (*TWF154, TWF132, TWF102*). To do this, we collected single conidia from each fungal strain and stained them directly with SR2200. For each of the three *A. oligospora* strains, we present in Fig. 4 (D, E and F) the last image of a temporal series obtained by FFT representing growth from a single conidium after 72 hours. FFT very accurately detected the mycelium for all strains (green networks in Figs. 4D, E and F). Moreover, the area covered by the mycelium (blue polygons) and the detected hyphal tips (yellow dots) were accurately characterized in all strains. Averages of the different measures computed for six replicates of each strain and time-point are also presented as graphs in Fig. 4 (G, H and I). These graphs show that TWF154 grew faster and formed more hyphal tips and segments than the other two *A. oligospora* strains, with TWF154 having 4-fold more hyphal tips and segments than TWF102 by the end of the experiment (Fig. 4G and 4H). Furthermore, TWF154 colonized the entire studied growth area (Fig. 4I) within the 72-hour experimental period. We performed a Mann Whitney test (MWT) (28) to find statistical differences between pairs of growth curves. While the two first measures showed a significant difference between the studied strains, the area covered by the mycelium follows a similar trend for TWF102 and TWF132 (p-value 0.18, Supplementary Table S1). Thus, the mycelial networks formed by these strains have different densities. This finding would have been overlooked simply using colony radius to measure mycelial development, highlighting the need to compute several measurements in order to fully characterize mycelial dynamics.

We conducted a second experiment in which we added dye to different media instead of staining the spores. To do so, we placed a drop of spore solution obtained from three fungal species (*A. oligospora, T. reesei* and *N. crassa*) on plates of PDA (nutrient-rich) or LNM (nutrient-poor) media mixed with SR2200 and recorded the growth of the spores for 48 hours. In Figure 5 (A-F), we present images showing the changes in mycelial growth detected by FFT at time intervals for each fungal species and type of medium. These plots reveal clear differences in mycelial density and area coverage among the fungal species and for the different media, with these differences better reflected by the quantitative data produced by FFT and graphically presented in Figure 5 (G-L; averages of two replicates per strain, type of media, and time-point for the total number of hyphal tips, total mycelial length and the area covered by the mycelium). Sigmoidal growth curves were apparent for each of the three fungal species and for both types of media (Fig. 5, I and L), which is typical of filamentous fungal growth (29,30). A lag phase, the duration of which varied depending on the species, is followed by an almost exponential growth phase, after which most species reached a plateau (once the studied growth area was full of hyphae).

**Figure 5.**
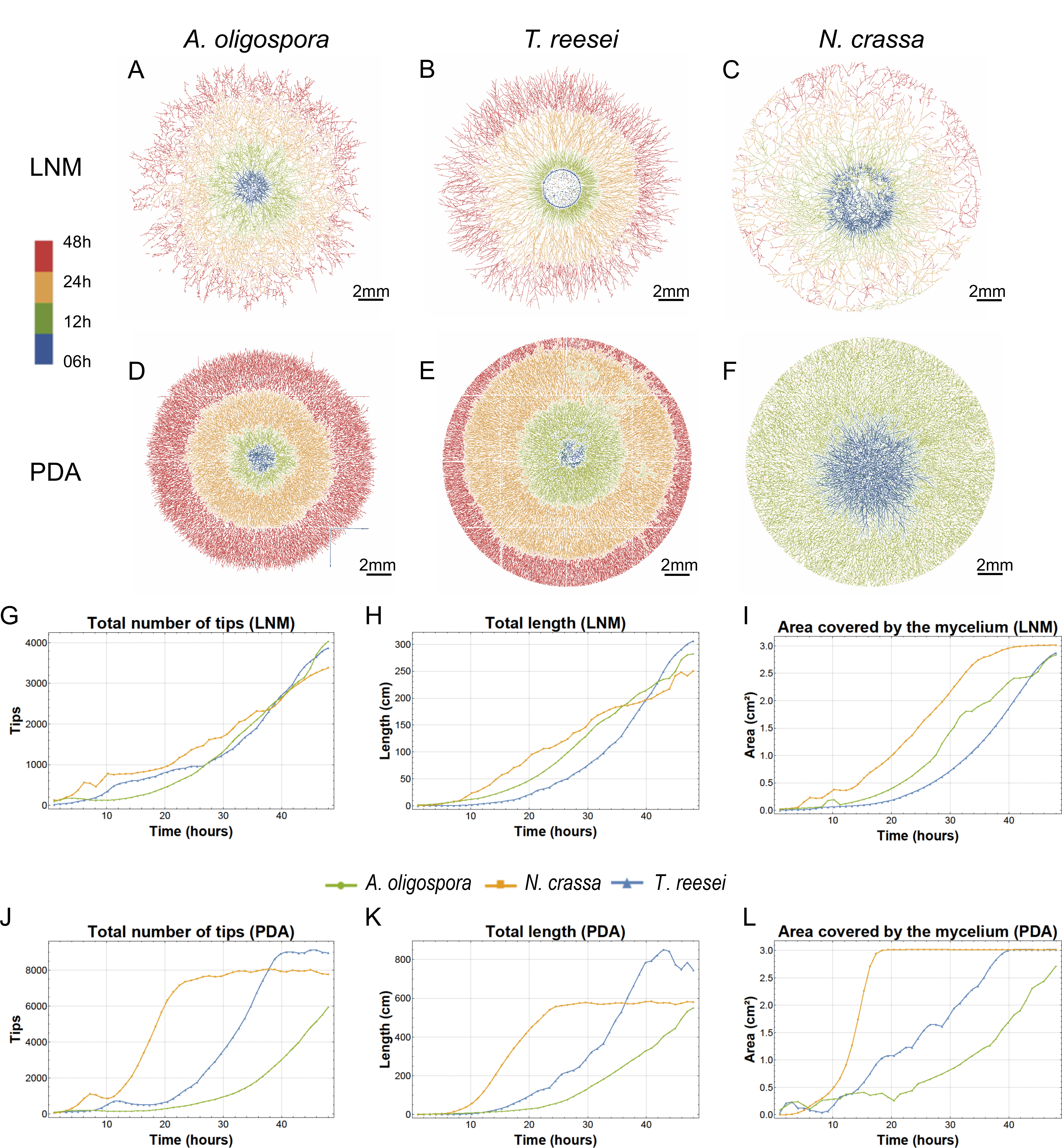
Mycelium development of different fungal species and in different media, as captured by FFT. Images showing the evolution of the mathematical graphs generated by FFT and representing 48 hours of growth on LNM (A-C) and PDA (D-F) media for each fungal species. Measures of mycelia over time, obtained as the mean of two replicates per species and time-point for LNM (G-I) and PDA (J-L) media.

Growth in the two different types of media differed considerably for each of the species, with all of them developing more than twice the number of hyphal tips and doubling their final total length in PDA compared to LNM. Interestingly, the area covered by mycelium is similar for both LNM and PDA by 48 hours, suggesting that all three species can efficiently colonize their surroundings irrespective of the media used. Differences in mycelial development were more pronounced in PDA in which each species exhibits very distinctive growth behavior resulting in low MWT p-values (Supplementary Table S2), especially compared to those obtained in LNM. Of the three species we assessed, *N. crassa* exhibited the most efficient colonizing behavior. This fungus had colonized the entire growth area in both nutrient conditions within 48 hours and, in PDA, it covered the full area within the first 24 hours of the experiment. However, *N. crassa* develops more sparse networks when compared to those of *T. reesei*. This latter fungus seems to develop more slowly, but generates more hyphal tips and a denser network. We found that *A. oligospora* grew more slowly than the other two species and did not cover the entire growth area within 48 hours. However, the final total mycelial length of *A. oligospora* was very similar to that of *N. crassa*, suggesting that *A. oligospora* makes more compact and dense networks.

### FFT can characterize complex fungal structures such as fungal traps

NTF develop traps to capture and consume nematodes. However, the trapping abilities of these fungi vary greatly, even among closely-related fungal species and in different types of media (4,31,32). Quantifying trap formation by different strains is crucial to study the trapping behavior of these fungal predators (4).

In order to quantify differences in trapping ability, we developed a trap counting function as part of the FFT program. FFT can very precisely distinguish traps from the rest of the mycelium (Fig. 6 A-C). In addition, FFT revealed significant differences in trap formation between the three assessed strains of *A. oligospora* (Fig. 6D). Each of the images in Figure 6 (A-C) represents an area of about 2.5 mm x 2.1 mm, and the average number of traps in an image ranged from 15 (TWF102) to 48 (TWF132).

**Figure 6.**
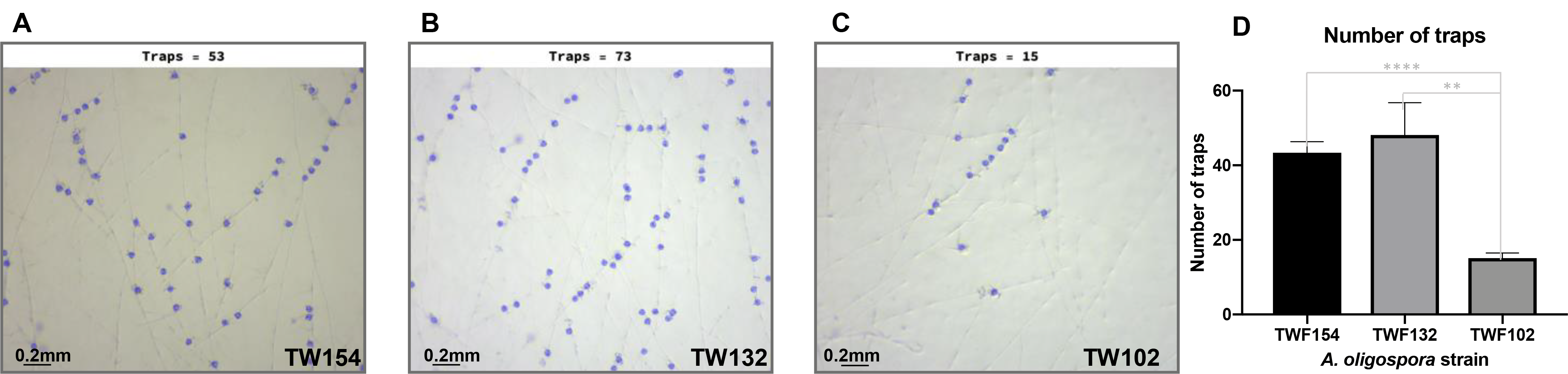
Performance of the trap counting function of FFT for three strains of *A. oligospora*. Images show the traps detected by FFT, highlighted in blue in the original images. The bar chart shows the difference in trap number between the different strains of *A. oligospora*, as computed using the average of three images representing the same replicate for a total of five replicates.

## Discussion

We have developed FFT, software for efficiently and quantitatively characterizing fungal growth and morphology. Through image analysis algorithms, FFT overcomes the subjectivity characteristic of manual analytical techniques and can simultaneously assess different phenotypic attributes. In addition, the complexity of the mycelium and morphological changes over time can be established using FTT. Moreover, FFT can examine several images at once, allowing semi-automatic analysis and giving rise to vast amounts of data capturing the biological diversity of fungi.

One of the main advantages of FFT is its versatility. Here, we used FFT to assess several phenotypic attributes from images captured with different imaging devices, at varying magnifications and resolutions. In addition, FFT utilizes simple image analysis functions to generate accurate results with minimal user interaction. For instance, only four parameters are needed to execute the entire FFT pipeline, and the effect of these parameters on the images can be examined in real time during the calibration process. Furthermore, apart from providing quantitative data outputs, FFT generates graphical outputs that facilitate visual inspection of results.

We focused on simple and easily available experimental techniques and imaging devices. All the experimental set-ups used in this manuscript are highly replicable, so most fungal research labs can generate results of similar quality. Other techniques and imaging devices may also produce images compatible with the different functions of FFT. However, certain criteria for image acquisition should be met to achieve optimal results with FFT. First, high contrast between the fungal features to be analyzed and the background is strongly recommended to aid the calibration process. Second, good image resolution is also required for FFT to produce reliable results. For example, a fungal spore or fungal trap should be represented by several pixels in order to be successfully characterized. Third, it is important to consider that image analysis may not be the best method to analyze structures occurring in three-dimensional space, such as aerial hyphae.

The scenarios studied in this manuscript only represent a small sample of all the possibilities that FFT has to offer. We focused on our own research interests and two model filamentous fungi (*N. crassa* and *T. reesei*), but the live-imaging techniques and image analysis algorithms described herein should produce analogous results for many filamentous fungi. Moreover, by using SR2200 to stain the fungal wall, we could follow fungal development without needing to perform genetic manipulations such as expressing GFP fluorescence, which can be challenging for many non-model fungal species (33). Considering the results obtained for the fungal strains and species we studied, FFT could be used to examine morphological differences in fungal growth and sporulation under different environmental conditions and between different fungal species. Furthermore, FFT can aid the study of genetic differences affecting the growth and morphology of fungi. For instance, one of the main obstacles to using NTF as biocontrol agents arises from their inability to become established in agricultural soils (6). FFT could be used to select fungal strains exhibiting enhanced trapping and colonization ability, potentially identifying strains that are more suited to be used as a biocontrol agents.

We wanted to provide the fungal research community with a simple tool for studying images of fungi at different stages of development. Image analysis tools that have been developed for other biological systems such as neurons (34), plant roots (35) and cells (36) may provide adequate results from fungal images. However, since fungi grow at a different scale, under different conditions and may be imaged using different devices to those systems, fungal imaging presents its own challenges, which is a major reason we decided to develop FFT specifically for the analysis of fungal images.

Unfortunately the wide scope and versatility of FFT hinders full automation of the image analysis process. In order to automatically calibrate the FFT algorithm, prior knowledge about the images is required, including the image acquisition settings and some morphological aspects of the studied fungi. The accuracy of the results generated by FFT depends on the calibration step, so adequate calibration is crucial. In addition, users should visually assess the quality of FFT outputs. For a more accurate performance, we suggest constructing ground truths in the images of interest and then computing image analysis performance measures (such as those described in the Supplementary Materials).

Overall, FFT represents a user-friendly tool that has been custom built for the analysis of fungal morphology and growth. It is highly versatile, allowing for the study of numerous fungal species and growth scenarios, and relies on images that can be generated from a wide range of devices. Therefore, we believe that incorporation of FFT into mycological laboratories will enhance the quantity and quality of data and present new avenues for fungal research.

## Materials and methods

### Organisms and media

Five filamentous fungal species: *Arthrobotrys oligospora* (*TWF154, TWF132 and TWF102*), *Arthrobotrys musifortis* (*TWF105*), *Arthrobotrys thaumasia*(*TWF678*), *Neurospora crassa (FGSC2489)* and *Trichoderma reseei* (*QM6a*) were studied in order to demonstrate the versatility of FFT. *Arthrobotrys* species are NTF able to capture and consume nematodes by developing complex adhesive networks (3). *N. crassa* and *T. reesei* were selected because they represent widely studied filamentous fungi (37,38). All fungal strains and species were cultured on Potato Dextrose Agar (PDA) medium plates at 25 **°**C in a dark incubator for 7 days prior to conducting experiments. In addition, *Caenorhabditis elegans* nematodes (N2) were used to trigger trap formation in *A. oligospora.*

### Image acquisition

#### Conidiation

Spore suspension of fungal strains was inoculated on 5 cm PDA (YPD for *N. crassa*) plates and incubated for 5 days at 25**°**C in the dark. 2 ml of ddH_2_O was added to each plate and conidia were carefully scratched off from the surface of the mycelium. The ddH_2_O from the plate was first filtered through two layers of non-woven cloth (to exclude extraneous hyphae) and then transferred to a 2 ml centrifuge tube for centrifugation at 13,000 x *g* for 1 minute. 1 ml of supernatant was extracted from each tube and 10 × 5 μl droplets of the remaining solution were placed on an LNM Petri-dish. Each droplet was imaged using a Zeiss Stemi 305 Stereo Microscope (Zeiss, Göttingen, Germany) and a Zeiss Axiocam ERc 5s Microscope Camera (Zeiss, Göttingen, Germany) at a resolution of 80X and 16X, for the conidiation study of fungal species and *A. oligospora* strains, respectively. The images comprised 2290×1920 pixels, representing an area of about 1.3 × 1 mm for the 80X images and 6.3 × 5.3 mm for the 16X images.

#### Conidia morphology

To study the morphology of conidia, conidia of different fungal species were stained by calcofluor white (Sigma) and imaged by an Axiovert 200M fluorescence microscope (Zeiss, Göttingen, Germany) at 400X using ultraviolet light and a blue/cyan filter. The resulting images represent an area of 347 × 260 μm and contain 1-3 conidia.

#### Trap counting

*A. oligospora* only develops traps in low nutrient environments in the presence of nematodes (4). Consequently, *A. oligospora* strains were first cultured for 2 days in 3.5 cm diameter LNM-containing Petri-dishes. Then, 30 adult living *C. elegans* nematodes were placed on the Petri-dishes for 6 hour and subsequently washed out from the plates with M9 buffer (22 mM KH_2_PO_4_, 42 mM Na2 HPO_4_, 86 mM NaCl). After an additional 18 hours of growth, *A. oligospora* strains were imaged using a Zeiss Stemi 305 Stereo Microscope (Zeiss, Göttingen, Germany) and a Zeiss Axiocam ERc 5s Microscope Camera (Zeiss, Göttingen, Germany). Three 40X images of the mycelium (size 2.5 × 2.1 mm) per plate were captured at random for 6 plates per strain, yielding a total of 18 images per strain.

#### Mycelium quantification

To characterize early mycelial growth of the *A. oligospora* strains, first single conidium were transferred to a 24 well plate containing a thin layer of LNM (200 μl). After 2 hours incubation, 1 ml of SCRI Renaissance 2200 (Renaissance Chemicals) dye (0.1%) was added to each well and led to act for 10 minutes in the dark. Then, each well was washed twice with PBS and the plates were placed on an ImageXpressMicro-XL system (Molecular Devices, Sunnyvale, California) set up at 25 °C and under controlled humidity. 12 images (1598 × 1598 pixels, size 0.66 × 0.66 cm) representing different parts of the well were captured at 2X magnification every hour for 72 hours. For each time point and well, the 12 images were assembled together (size 2.64 × 2.64 cm) and a mask representing the edge of the well was subtracted.

The growth of the fungal species in different growth media was assessed by adding 7.5μl of SR2200 dye (0.05%) to 15 ml of each type of media (LNM and PDA). 300μl of the stained media was placed in a 12 well plate and 5μl of conidia solution was inoculated on two wells per species and media. ImageXpressMicro-XL system (Molecular Devices, Sunnyvale, California) was used to capture 2X images every hour for 48 hours. Finally, the images representing each well per time point were assembled resulting in images of 6392 × 6392 pixels and a mask representing the well edge was subtracted from them.

#### Fungal Feature Tracker (FFT) interface and workflow

The general workflow of FFT consists of the following three consecutive steps: input selection, calibration, and output configuration. For simplicity, this workflow is common to all quantification functions of FFT, i.e., the workflow is the same irrespective of the phenotypic feature being analyzed. In addition, the user interface is divided into simple tabs, each representing one of the workflow steps (Fig. 7), thereby ensuring execution of FFT is user-friendly.

**Figure 7.**
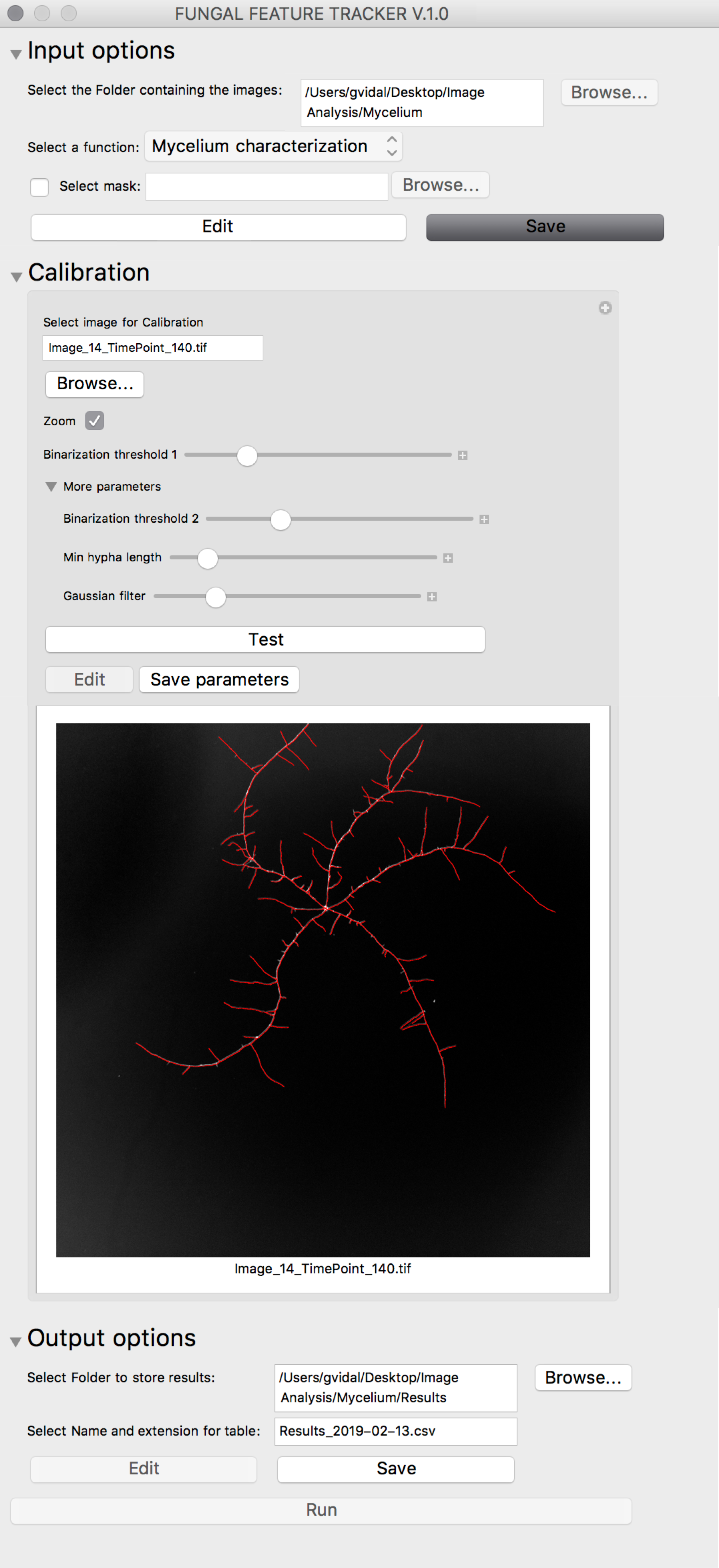
General workflow of FFT. The “Input options” tab (Top) allows users to select the set of images for analysis and the quantification function to be applied. In the “Calibration” tab (Middle), users can select and test the effect of different parameter combinations on one image from the set of images. FFT output information is defined in the “Output options” tab (Bottom). Once all this information has been provided, the “Run” button executes FFT analysis on all images in the set of images.

The first tab is “Input options”. In this tab, users can select a folder containing the set of images to be analyzed by FFT. Importantly, all images in the selected folder will be analyzed using the same parameters and, therefore, they should be as homogeneous as possible. For instance, images obtained with different devices or using very different settings should not be part of the same set of images. Additionally, a mask can be selected in this tab to reduce noise present in the set of images. For example, in our case, masks representing the edge of the wells were subtracted from the original images. Since the same mask is subtracted from all images of a given set, we strongly advise only using simple and general masks. Also in the “Input options” tab, the quantification function must be selected. There are four available functions in the current version of FFT: conidia/spore counting, conidia/spore morphology characterization, trap counting, and mycelium characterization. Each of these functions rely on different image analysis algorithms, have their own specific parameters, and result in different outputs. In the section below, we explain the different quantification functions of FFT in more detail.

Once the set of images and the FFT function have been selected, the “Calibration” tab is enabled. The purpose of this tab is to select the parameters that work best with all the images of the set and the selected function. Values of the different parameters can be changed in this tab using sliders. When the value of a parameter is changed, a screen at the center of the “Calibration” tab is updated to show how the chosen parameters affect the output of the function. By default, the output of the selected function is shown for an image randomly selected from among those in the image set, but users can select any image on which to perform the calibration. The outcome of applying the chosen parameters on several images of the set can be checked by clicking the “Test” button. The “Test” button opens a new window showing the output obtained from applying the selected parameters to three randomly selected images of the set. Once an optimal parameter combination has been selected, users must save these parameters by clicking “Save parameters”. The calibration can also be performed on a augmented area of an image by selecting the “Zoom” checkbox. This feature is particularly useful for calibrating large images in which features cannot be easily observed using the whole image and to speed up the calibration process since small images can be analyzed faster by FFT.

The final tab, “Output options”, allows users to select the location where the FFT outputs are stored. Upon choosing the output directory, the “Run” button becomes active. By clicking this button, the function selected in the “Input options” tab is automatically applied according to the parameters chosen in the “Calibration step” to all images of the selected set. During this execution phase, several files representing the outputs of FFT are generated and stored in the output folder selected in the “Output options” tab.

FFT produces three different outputs irrespective of the selected function. First, a configuration .txt file containing the values of the parameters applied during the execution phase. Second, a table of results in which the first column corresponds to the name of the image and the remaining columns represent features extracted from the image. The extension and name of this table of results can be modified in the “Output options” tab. Third, FFT generates an output image for each image of the set, which represents the original image overlaid with the detected features.

The complete source code of FFT, together with more information on how to execute it and some test images of fungi, can be found at the Hsueh lab github: https://github.com/hsueh-lab/FFT.

### FFT functions

#### Conidia/spore counting

The purpose of the conidia/spore counting function is to detect and count the number of spores or conidia spread over a transparent background. This function relies on the fact that conidia or spores are darker than the surrounding background. Therefore, strong contrast between the spores and the background in the original images is crucial for this function to perform satisfactorily. In the first step, the original image is transformed into a binary (black and white) image containing only the most important information. To do this, we first inverted the colors of the image so that the spores are represented as white objects against a dark background. We used the `MorphologicalBinarize` function from Mathematica (Version 11.1.1.0, Wolfram Research Inc., USA) to generate the binary image. This function requires selection of two parameters (the binarization thresholds b1 and b2) in FFT during the calibration step. All image pixels having value greater than the selected threshold for b1 are assigned a value of 1 (white). In addition, all pixels whose value is above the selected b2 threshold and that are connected to the foreground are also assigned a value of 1. The resulting binary image contains several objects in white against a black background. Some of these objects might be spores/conidia and some may represent noise or contamination (such as hyphae or dust). Consequently, users must determine which objects are spores/conidia by filtering according to morphology. To do this, minimum and maximum areas for spores can be defined, as can an elongation threshold. These parameters can be applied to filter out tiny particles that might represent dust or large, long and narrow artifacts that may correspond to hyphal fragments. These three thresholds can be set in the “Calibration” tab, where they are referred to as “Min Area”, “Max Area” and “Elongation”, respectively. To select the objects that pass this filter, i.e., spores/conidia, we used the `SelectedComponents` function from Mathematica with the abovementioned parameters. The detected spores/conidia are then highlighted in blue and overlaid on the original image as an output image (Fig. 2 and 3). Finally, the detected spores/conidia of each image of the set are counted to generate the results table.

#### Conidia/spore morphology

The algorithm of the conidia/spore morphology characterization function is very similar to that used for conidia/spore counting. As for the previous function, binary images are generated using the `MorphologicalBinarize` function. However, in this case, the spores/conidia have been treated with calcofluor white so it is not necessary to invert the colors since the spores already appear white against a dark background. This function also uses `SelectedComponents` to detect and filter the forms representing spores/conidia based on thresholds for minimum and maximum areas and elongation. Thus, the parameters used for the conidia/spore morphology function and the conidia/spore counting function are the same: the binarization thresholds “b1” and “b2”, and the spore morphology parameters “Min Area”, “Max Area” and “Elongation”. It must be noted that due to differences in the scales of the input images, the thresholds for minimum and maximum area, as well as the elongation parameter, can differ considerably for these two functions. For each image, the output of the conidia/spore morphology function consists of the area (in pixels) of each conidium present in the image, its length, width and circularity. Here, we define length as the longest axis of the best-fit ellipse fitted to the detected spore/conidia, whereas width corresponds to the shortest axis of the same ellipse. Circularity is defined as the ratio of the equivalent disk perimeter to the actual perimeter length, so a circular spore has a circularity value close to 1. Values for these features are presented in the output images, graphically overlaid on the original image (as shown in Fig. 1), and detailed for each image in the results table.

#### Mycelium characterization

The image analysis algorithm behind the mycelium characterization function is the most complex employed by FFT and it is adapted from previously published algorithms (29,39). The first step of this algorithm is to simplify the information contained in the raw images by reducing noise. To do this, we applied a Gaussian filter (using the `GaussianFilter` function from Mathematica) to the original image to create a mask that is then subtracted from the original image. This function uses a Gaussian kernel of a given radius (in pixels) to convolve the image, i.e., it replaces each pixel by a linear combination of its neighboring pixels according to the weights given by the Gaussian kernel. This procedure results in a blurry image that reduces noise, homogenizes the background, and accentuates the most prominent pixels in the image upon being subtracted from the original image. The radius of the Gaussian kernel can be selected by the user in the “Calibration” tab under the name “gaussian”. The noise-reduced image is then transformed into a binary image using the `MorphologicalBinarize` function defined by the binarization thresholds b1 and b2. We often observed gaps in this binary image due to out-of-focus areas and areas of low contrast due to uneven distribution of the dye (especially at the edges of Petri-dishes). To overcome this issue, we performed an image dilation using a 3×3 box matrix via the `MorphologicalTransform` function of Mathematica. As shown in Figure 8B, this step results in a binary image showing thick hyphae and a connected mycelium. After this dilation step, we used the `Thinning` function of Mathematica to transform the thick hyphae into lines of the same width. Finally, we deleted small components and pruned small uninformative branches (see Figure 8C) using the functions `DeleteSmallComponents` and `Pruning`, respectively, with the parameter “Minimum hyphae” that can be selected during the calibration step. In addition, if users define a mask, it can be subtracted from the final binary image to delete misleading information such as Petri-dish edges or other evident noise arising from the experimental set-up.

**Figure 8.**
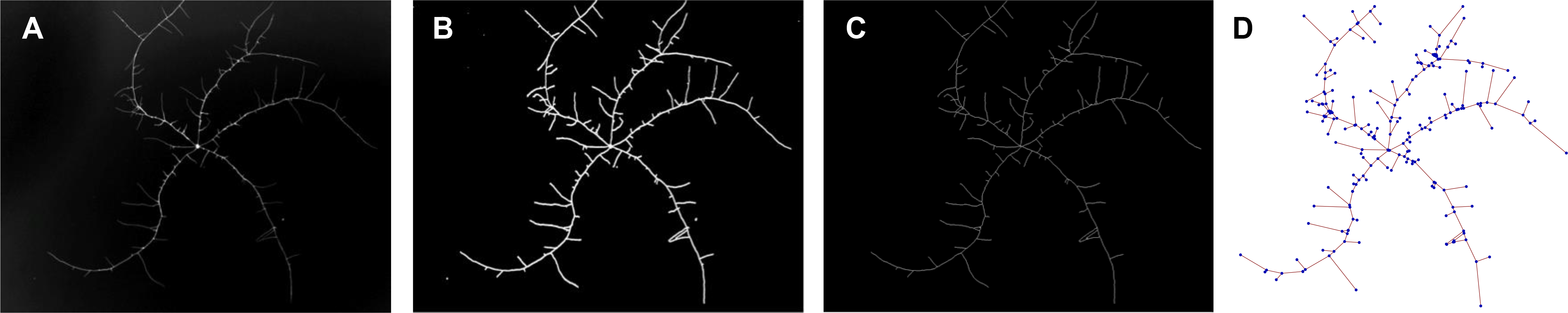
Workflow of the mycelium characterization algorithm. Original image (A). Binary image obtained by subtracting a gaussian-filtered version of the original image, applying morphological binarization, and applying a maximum area filter (B). Result of applying the `Thinning` function to the image in B (C). Mathematical graph obtained using the `MorphologicalGraph` function, in which nodes (blue) represent hyphal junctions/tips and edges (red) represent hyphae (D).

Once a clear, binary image has been obtained in which the hyphae in the mycelium are shown as simple lines, it is transformed into a mathematical graph in which nodes represent objects and edges represent relationships between the objects (40). In our case, each node represents a hyphal tip or junction and each edge represents the hyphae connecting those junctions (41). We created mathematical graphs using the `MorphologicalGraph` function of Mathematica. Several features can be quantified from these graphs representing the mycelium. For instance, we obtained growth-related measures such as the total length of the mycelium (as the sum of all the edges in the graphs (42)), the number of hyphal tips (as the nodes connected to only one edge), and the area of the mycelium (as the convex hull of the nodes composing the graph). We refer the reader to (29) for more detail about these measures.

The final output of the mycelium characterization function consists of a table of results in which each row represents an image together with the respective values for all of the aforementioned features. In addition, output images are generated for each original image that display the mathematical graph, the detected hyphal tips and the computed area of the mycelium highlighted over the original image. For instance, in Fig. 4, we present output images obtained using the mycelium characterization function in which the area covered by the mycelium is shown in blue, the detected hyphae are shown in green, and the hyphal tips are shown in yellow. It is worth noting that even though the mycelium characterization function can be used on single images, it can also be used to study fungal development over time. For instance, by using a temporal series of images as the image set, it is possible to characterize mycelium features at consecutive time-points and consequently to track the changes occurring in the mycelium over time.

#### Trap counting

The trap counting function of FFT was designed to count adhesive network traps; a feature observed only in certain NTF (4). However, we believe that other fungal traps such as constraining rings (43) and other complex structures such as appressoria (44) could also be recognized by using this function since the traps are detected as structures distinct from the rest of the mycelium. Therefore, as for the previously described functions, a binary image must be generated and traps can then be selected based on their morphology, i.e., their size and elongation. Thus, the parameters applied in this function do not differ from those of the first two functions, so the trap counting function could be considered a particular application of the conidia/spore counting function.

## Acknowledgments

The authors thank Ting-Fang Wang at IMB Academia Sinica for sharing the *Neurospora* and *Trichoderma* strains, and Pedro Gonçalves for helpful comments on the manuscript. We also thank all the members of the Hsueh lab for their help and support. This work was supported by the Academia Sinica Postdoctoral Fellowship to GV and the start-up fund of Academia Sinica and Ministry of Science and Technology 106-2311-B-001-039-MY3 to YPH.

## Figure captions supplementary materials

**Figure S1**. Comparison of conidial morphology for three *A. oligospora* strains. Spore area (A), length (B), width (C) and circularity (D), as computed by FFT using a total of 10 images per fungal strain.

**Figure S2**. Comparison of an original image, ground truth image, and the mycelium detected by FFT for two images of *A. oligospora*. Original image 9 following mask subtraction (A), ground truth computed manually from the original image (B), and the image obtained by FFT after applying filters and binarization (C). Original image (D), ground truth image (E) and FFT-detected image (F) from a zoomed in view of image 4 showing an early-stage mycelium of *A. oligospora.*

## Notes

https://github.com/hsueh-lab/FFT

## Bibliography

1. Boddy L. Saprotrophic Cord-Forming Fungi: Meeting the Challenge of Heterogeneous Environments. Mycologia [Internet]. 1999 Jan;91(1):13. Available from: http://www.jstor.org/stable/10.2307/3761190?origin=crossref

2. Magan N. 6 Fungi in Extreme Environments. Environ Microb Relationships Environ Microb relationshipsIV. 2007;4:85.

3. Nordbring-Hertz B, Jansson H-B, Tunlid A. Nematophagous Fungi. In: eLS [Internet]. Chichester, UK: John Wiley & Sons, Ltd; 2011. p. 1–13. Available from: 10.1002/9780470015902.a0000374.pub3

4. Vidal-Diez de Ulzurrun G, Hsueh Y. Predator-prey interactions of nematode-trapping fungi and nematodes⍰: both sides of the coin. 2018;3939–49. Available from: https://doi.org/10.1007/s00253-018-8897-5

5. Lopez-Llorca L V., Maciá-Vicente JG, Jansson H-B. Mode of Action and Interactions of Nematophagous Fungi. In: Ciancio A, Mukerji KG, editors. Integrated Management and Biocontrol of Vegetable and Grain Crops Nematodes. Dordrecht: Springer Netherlands; 2007. p. 51–76. (Integrated Management of Plant Pests and Diseases; vol. 2).

6. Jaffee BA. Correlations between most probable number and activity of nematode-trapping fungi. Phytopathol 93, 1599–1605. 2003;93(12):1599–605.

7. Krull R, Wucherpfennig T, Esfandabadi ME, Walisko R, Melzer G, Hempel DC, et al. Characterization and control of fungal morphology for improved production performance in biotechnology. J Biotechnol. 2013;163(2):112–23.

8. Meyer V. Genetic engineering of filamentous fungi - Progress, obstacles and future trends. Biotechnol Adv. 2008;26(2):177–85.

9. Pasanen A-L, Kalliokoski P, Pasanen P, Jantunen MJ, Nevalainen A. Laboratory studies on the relationship between fungal growth and atmospheric temperature and humidity. Environ Int. 1991;17(4):225–8.

10. Song T-Y, Xu Z-F, Chen Y-H, Ding Q-Y, Sun Y-R, Miao Y, et al. Potent Nematicidal Activity and New Hybrid Metabolite Production by Disruption of a Cytochrome P450 Gene Involved in the Biosynthesis of Morphological Regulatory Arthrosporols in Nematode-Trapping Fungus Arthrobotrys oligospora. J Agric Food Chem [Internet]. 2017;65(20):4111–20. Available from: http://pubs.acs.org/doi/10.1021/acs.jafc.7b01290

11. Sun M, Liu X. Carbon requirements of some nematophagous, entomopathogenic and mycoparasitic Hyphomycetes as fungal biocontrol agents. Mycopathologia. 2006;161(5):295–305.

12. Ajitomi A, Taba S, Ajitomi Y, Kinjo M, Sekine K. Efficacy of a Simple Formulation Composed of Nematode-Trapping Fungi and Bidens pilosa var. radiata Scherff Aqueous Extracts (BPE) for Controlling the Southern Root-Knot Nematode. Microbes Environ. 2017;33(1):4–9.

13. Li DP, Holdom DG. Effects of Nutrients on Colony Formation, Growth, and Sporulation of Metarhizium anisopliae .pdf. Journal of Invertebrate Pathology. 1995.

14. Sherif SM, Shukla MR, Murch SJ, Bernier L, Saxena PK. Simultaneous induction of jasmonic acid and disease-responsive genes signifies tolerance of American elm to Dutch elm disease. Sci Rep [Internet]. 2016;6(February):21934. Available from: http://www.nature.com/srep/2016/160223/srep21934/full/srep21934.html

15. Xie L, Han JH, Kim JJ, Lee SY. Effects of culture conditions on conidial production of the sweet potato whitefly pathogenic fungus Isaria javanica. Mycoscience [Internet]. 2016;57(1):64–70. Available from: http://dx.doi.org/10.1016/j.myc.2015.09.002

16. Singh UB, Sahu A, Sahu N, Singh RK, Renu S, Singh DP, et al. Arthrobotrys oligosporamediated biological control of diseases of tomato (Lycopersicon esculentum Mill.) caused by Meloidogyne incognita and Rhizoctonia solani. J Appl Microbiol. 2013;114(1):196–208.

17. Seiler S, Plamann M. The Genetic Basis of Cellular Morphogenesis in the Filamentous Fungus Neurospora crassa. Mol Biol Cell [Internet]. 2003 Nov;14(11):4352–64. Available from: http://www.molbiolcell.org/doi/10.1091/mbc.e02-07-0433

18. Bailey-Shrode L, Ebbole DJ. The fluffy Gene of Neurospora crassa Is Necessary and Sufficient to Induce Conidiophore Development. Genetics. 2004;166(4):1741–9.

19. Papagianni M. Characterization of Fungal Morphology using Digital Image Analysis Techniques. J Microb Biochem Technol. 2014;06(04):189–94.

20. Brunk M, Sputh S, Doose S, Van De Linde S, Terpitz U. HyphaTracker: An ImageJ toolbox for time-resolved analysis of spore germination in filamentous fungi. Sci Rep. 2018;8(1):1–13.

21. Benyon FHL, Jones AS, Tovey ER, Stone G. Differentiation of allergenic fungal spores by image analysis, with application to aerobiological counts. Aerobiologia (Bologna). 1999;15(3):211–23.

22. Tronnolone H, Gardner JM, Sundstrom JF, Jiranek V, Oliver SG, Binder BJ. TAMMiCol: Tool for analysis of the morphology of microbial colonies. PLOS Comput Biol. 2018;14(12):1–15.

23. Lecault V, Patel N, Thibault J. Morphological Characterization and Viability Assessment of Trichoderma reesei by Image Analysis. Biotechnol Prog [Internet]. 2008 Sep 5;23(3):734–40. Available from: http://doi.wiley.com/10.1021/bp0602956

24. Kessel GJT, De Haas BH, der Plas CH, Meijer EMJ, Dewey FM, Goudriaan J, et al. Quantification of mycelium of Botrytis spp. and the antagonist Ulocladium atrum in necrotic leaf tissue of cyclamen and lily by fluorescence microscopy and image analysis. Phytopathology. 1999;89(10):868–76.

25. Mycologia S, Jun NM. Two New Fluorescent Dyes Applicable for Visualization of Fungal Cell Walls Author (s): H.C. Hoch, C.D. Galvani, D.H. Szarowski and J.N. Turner Stable URL : http://www.jstor.org/stable/3762339 REFERENCES Linked references are available on J. 2017;97(3):580–8.

26. 70. Harris K, Crabb D, Young IM, Weaver H, Gilligan CA, Otten W, et al. In situ visualisation of fungi in soil thin sections: Problems with crystallisation of the fluorochrome FB 28 (Calcofluor M2R) and improved staining by SCRI Renaissance 2200. Mycol Res. 2002;106(3):293–7.

27. Wang G, Lopez-Molina C, Vidal-Diez de Ulzurrun G, De Baets B. Noise-robust line detection using normalized and adaptive second-order anisotropic Gaussian kernels. Signal Processing [Internet]. 2019;160:252–62. Available from: https://doi.org/10.1016/j.sigpro.2019.02.027

28. Mann HB, Whitney DR. On a Test of Whether one of Two Random Variables is Stochastically Larger than the Other. Ann Math Stat [Internet]. 1947 Mar;18(1):50–60. Available from: http://www.scirp.org/journal/doi.aspx?DOI=10.4236/tel.2012.21009

29. Vidal-Diez de Ulzurrun G, Baetens JM, Van den Bulcke J, Lopez-Molina C, De Windt I, De Baets B. Automated image-based analysis of spatio-temporal fungal dynamics. Fungal Genet Biol [Internet]. 2015;84:12–25. Available from: http://dx.doi.org/10.1016/j.fgb.2015.09.004

30. Barry DJ, Williams GA. Microscopic characterisation of filamentous microbes: towards fully automated morphological quantification through image analysis. J Microsc. 2011;244(1):1–20.

31. Su H, Zhao Y, Zhou J, Feng H, Jiang D, Zhang KQ, et al. Trapping devices of nematodetrapping fungi: formation, evolution, and genomic perspectives. Biol Rev. 2017;92(1):357–68.

32. Saxena G, Dayal R, Mukerji KG. Interaction of nematodes with nematophagus fungi: induction of trap formation, attraction and detection of attractants. FEMS Microbiol Lett. 1987;45(6):319–27.

33. Lorang JM, Tuori RP, Martinez JP, Sawyer TL, Redman RS, Rollins JA, et al. Green Fluorescent Protein Is Lighting Up Fungal Biology. Appl Env Microbiol. 2001;67(5):1987–94.

34. Kim KM, Son K, Palmore GTR. Neuron image analyzer: Automated and accurate extraction of neuronal data from low quality images. Sci Rep [Internet]. 2015;5(November):1–12. Available from: http://dx.doi.org/10.1038/srep17062

35. Cai J, Zeng Z, Connor JN, Huang CY, Melino V, Kumar P, et al. RootGraph: A graphic optimization tool for automated image analysis of plant roots. J Exp Bot. 2015;66(21):6551–62.

36. O’Brien J, Hayder H, Peng C. Automated Quantification and Analysis of Cell Counting Procedures Using ImageJ Plugins. J Vis Exp [Internet]. 2016 Nov 17;(117). Available from: http://www.jove.com/video/54719/automated-quantification-analysis-cell-counting-procedures-using

37. Borkovich K, Alex L, Yarden O, Freitag M, Turner GE, Read ND, et al. Lessons from the Genome Analysis of Neurospora crassa. Microbiol Mol Biol Rev [Internet]. 2004;68(1):1–108. Available from: mmbr.asm.org

38. Martinez D, Berka RM, Henrissat B, Saloheimo M, Arvas M, Baker SE, et al. Genome sequencing and analysis of the biomass-degrading fungus Trichoderma reesei (syn. Hypocrea jecorina). Nat Biotechnol [Internet]. 2008 May 4;26(5):553–60. Available from: http://www.nature.com/articles/nbt1403

39. Lopez-Molina C, Vidal-Diez De Ulzurrun G, Baetens JM, Van Den Bulcke J, De Baets B. Unsupervised ridge detection using second order anisotropic Gaussian kernels. Signal Processing [Internet]. 2015;116:55–67. Available from: http://dx.doi.org/10.1016/j.sigpro.2015.03.024

40. Gibbons A. Alghoritmic graph theory. 1985.

41. Obara B, Grau V, Fricker MD. A bioimage informatics approach to automatically extract complex fungal networks. Bioinformatics. 2012;28(18):2374–81.

42. Trinci APJ. A Study of the Kinetics of Hyphal Extension and Branch Initiation of Fungal Mycelia. J Gen Microbiol [Internet]. 1974 Mar 1;81(1):225–36. Available from: http://mic.microbiologyresearch.org/content/journal/micro/10.1099/00221287-81-1-225

43. Liu K, Zhang W, Lai Y, Xiang M, Wang X, Zhang X, et al. Drechslerella stenobrocha genome illustrates the mechanism of constricting rings and the origin of nematode predation in fungi. BMC Genomics [Internet]. 2014;15(1):114. Available from: http://bmcgenomics.biomedcentral.com/articles/10.1186/1471-2164-15-114

44. Kong LA, Li GT, Liu Y, Liu MG, Zhang SJ, Yang J, et al. Differences between appressoria formed by germ tubes and appressorium-like structures developed by hyphal tips in Magnaporthe oryzae. Fungal Genet Biol [Internet]. 2013;56:33–41. Available from: http://dx.doi.org/10.1016/j.fgb.2013.03.006

