## Supplementary figures and images for "Fungal Feature Tracker (FFT): A tool for quantitatively characterizing the morphology and growth of filamentous fungi"

### Supplemental Figure1

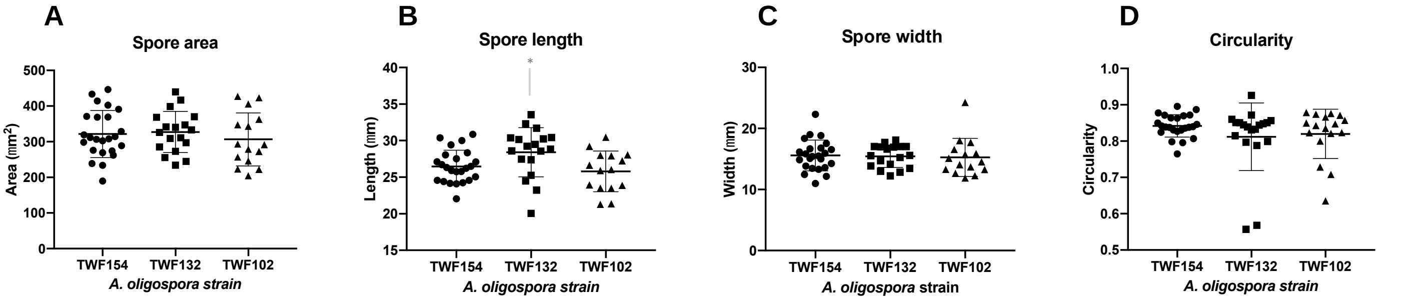

### Supplemental Figure2

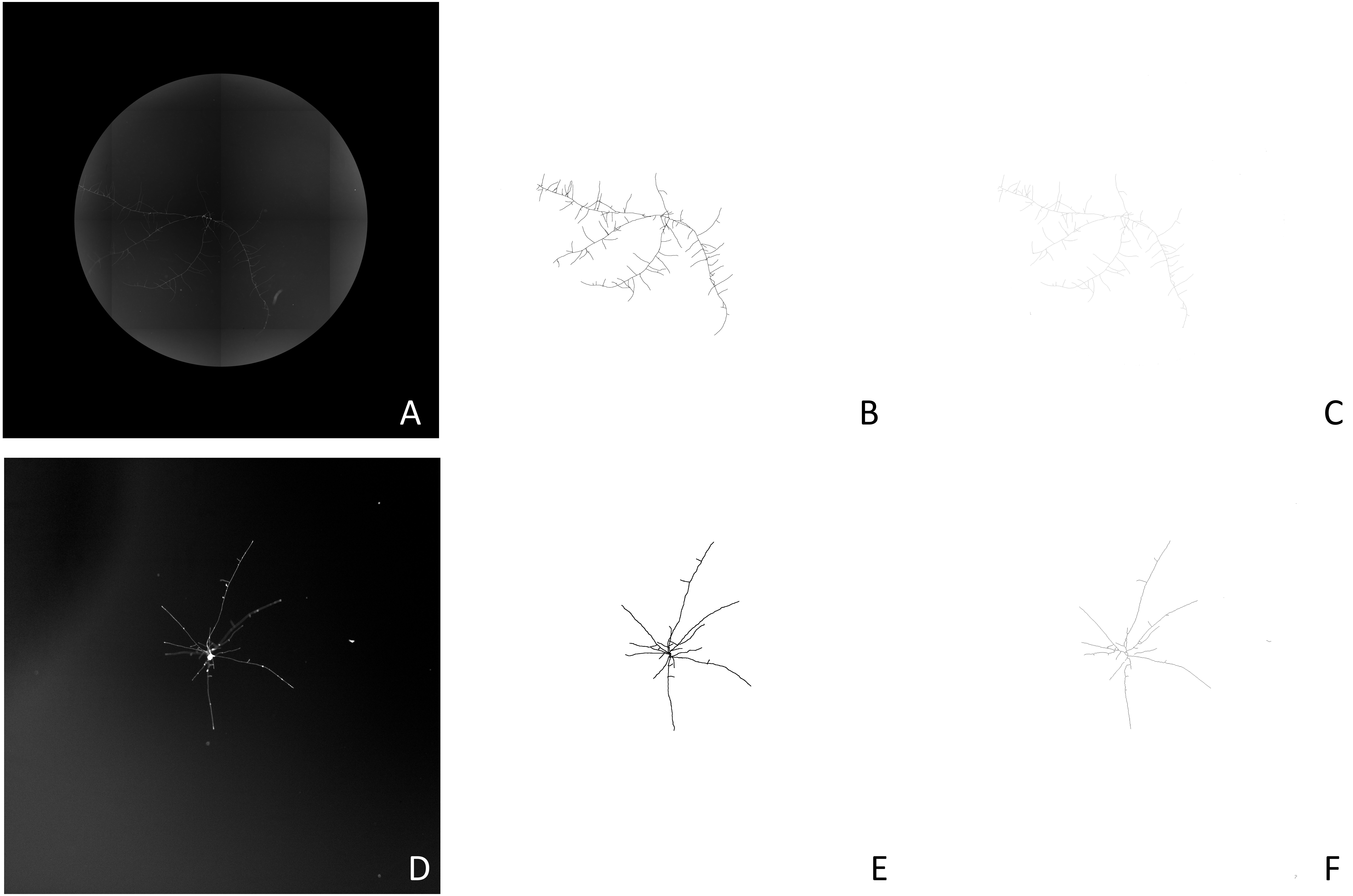
