## Supplemental information for "Fungal Feature Tracker (FFT): A tool for quantitatively characterizing the morphology and growth of filamentous fungi"

**Supplementary materials**

**Accuracy measurement of the edge detection algorithm applied in the mycelium characterization function of FFT**

We measured the accuracy of the fungal networks extracted by the mycelium characterization function of FFT using 10 images showing different stages of the mycelium developed by *A. oligospora* (strain TWF154). To do this, we first made ground truths by manually tracing the mycelium of each of the original images. We compared these ground truths with the images generated by FFT prior to conversion into mathematical graphs (see Figure S2). Since displacement is common in edge detection algorithms and particularly after applying a Thinning filter, we allowed for a 2-pixel displacement from the ground truth, i.e., a pixel in the FFT image that is positioned 2 pixels away from its real location was still considered successfully detected.

Using the aforementioned criteria, we computed the F-measure and Matthews Correlation Coefficient (MCC), both measures widely used to assess the performance of image detection algorithms (27). The F-measure captures the harmonic mean between precision (Prec) and recall (Rec) and is given by the formula:

$$F=\frac{2\cdot Prec\cdot Rec}{Prec+Rec}$$

with

$\mathrm{Prec}=\frac{TP}{TP+FP}$ and $Rec=\frac{TP}{TP+FN}$ ,

where TP, FP and FN are the number of true positives, false positives and false negatives, respectively. For each image, pixels detected by FFT and also present in the ground truth were considered true positives, pixels detected by FFT and not present in the ground truth were false positives, pixels not detected by FFT and not present in the ground truth were true negatives, and those not detected by FFT but present in the ground truth were false negatives.

The MMC can be used as an evaluation metric for computing the correlation between data detected by an image analysis algorithm and the ground truth. The MCC formula is given by:

$$\mathrm{MCC}=\frac{\left( TP\cdot TN \right)-(FP\cdot FN)}{\sqrt{(TP+FP)\cdot(TP+FN)\cdot(TN+FP)\cdot(TN+FN)}}$$

where TP, FP, TN, FN represent true positives, false positives and false negatives, respectively.

| **Fungal Strains** | **Total number of tips** | **Total length** | **Area covered by the mycelium** |
| --- | --- | --- | --- |
| TWF102 vs TWF132 | 0.0003 | 0.0009 | 0.18 |
| TWF102 vs TWF154 | 0.0081 | 0.0049 | 1.07E-06 |
| TWF132 vs TWF154 | 1.65E-08 | 6.30E-08 | 1.40E-09 |

**Table S1.** Mann Whitney test results from the growth comparison of *A. oligospora* strains. P-values obtain for each measure and pair of strains computed from the mean of six replicates per strain and time-point.

| **Species and Media** | **Total number of tips** | **Total length** | **Area covered by the mycelium** |
| --- | --- | --- | --- |
| T.R vs N.C LNM | 0.25 | 0.039 | 0.0005 |
| T.R vs A.O LNM | 0.5 | 0.112 | 0.1418 |
| N.C vs A.O LNM | 0.072 | 0.64 | 0.0185 |
| T.R vs N.C PDA | 0.0458 | 0.1562 | 0.0134 |
| T.R vs A.O PDA | 0.0075 | 0.1015 | 0.0181 |
| N.C vs A.O PDA | 2.85E-08 | 8.24E-06 | 2.25E-06 |

**Table S2.** Mann Whitney test results from the growth comparison of the different fungal species. P-values obtain for each measure, media condition and species combination computed from the mean of two replicates per species and time-point.
